# iRAP - an integrated RNA-seq Analysis Pipeline

**DOI:** 10.1101/005991

**Authors:** Nuno A. Fonseca, Robert Petryszak, John C. Marioni, Alvis Brazma

## Abstract

RNA-sequencing (RNA-Seq) has become the technology of choice for whole-transcriptome profiling. However, processing the millions of sequence reads generated requires considerable bioinformatics skills and computational resources. At each step of the processing pipeline many tools are available, each with specific advantages and disadvantages. While using a specific combination of tools might be desirable, integrating the different tools can be time consuming, often due to specificities in the formats of input/output files required by the different programs.

Here we present iRAP, an integrated RNA-seq analysis pipeline that allows the user to select and apply their preferred combination of existing tools for mapping reads, quantifying expression, testing for differential expression. iRAP also includes multiple tools for gene set enrichment analysis and generates web browsable reports of the results obtained in the different stages of the pipeline. Depending upon the application, iRAP can be used to quantify expression at the gene, exon or transcript level. iRAP is aimed at a broad group of users with basic bioinformatics training and requires little experience with the command line. Despite this, it also provides more advanced users with the ability to customise the options used by their chosen tools.

iRAP is available under General Public License 3 (GPLv3) and although it should be portable to any POSIX-compliant operating system, several third party programs only run on Linux. iRAP can be obtained from http://code.google.com/p/irap.

## 1 Introduction

A typical RNA-seq analysis pipeline involves several stages (Fig. 1), starting with processing of the raw reads and moving ultimately to identification of differentially expressed genes. At each step of the pipeline the user is faced with choosing one of many existing tools. However, efficiently integrating a chosen set of tools is typically not straightforward, due primarily to differences in the input file formats required by each tool. In an attempt to overcome this problem, several pipelines for analyzing RNA-seq data have been developed (see Table 1 in Appendix for a summary). Grape [10] exploits a single mapper with different mapping strategies before using FluxCapacitor for quantification. Array express HTS [6], an R based pipeline, performs the mapping using BWA [13], Bowtie [12] or Tophat [22] before using Cufflinks [23] or MMSeq [25] to quantify expression levels. Finally, Galaxy [5] is a general purpose web-based platform that allows users to analyse different types of HTS data - for RNA-seq, Galaxy supports the Tuxedo pipeline (Tophat, Cufflinks and Cuffdiff).

**Figure 1.**
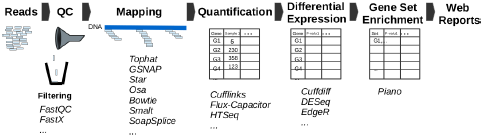
iRAP: Main steps in the analysis of RNA-seq data.

**Table 1:** iRAP comparison to other RNA-seq analysis pipelines. QC/Filter: indicates whether pipelines (can) apply quality control / allow reads to be filtered. Mappers: the number of mappers supported; Quantification: the number of quantification methods supported. DE: the number of differential expression analysis methods supported. GSE: whether the pipeline integrates gene set enrichment analysis. Web reports: indicates if pipelines produce a web report for each stage of the analysis. *Mappers part of the RNA-seq analysis in Galaxy.

| Pipeline | QC/Filter | Mappers | Quantification | DE | GSE | Web Reports |
| --- | --- | --- | --- | --- | --- | --- |
| Galaxy (RNA-seq) [5] | Yes/Yes | 1* | 1 | 1 | No | No |
| ArrayExpress HTS [6] | Yes/No | 3 | 2 | No | No | Yes |
| Grape [10] | Yes/No | 1 | 1 | No | No | Yes |
| PRADA [20] | No/No | 1 | 1 | No | No | No |
| RSeqFlow [27] | No/No | 2 | 1 | 2 | No | No |
| iRAP | Yes/Yes | 11 | 5 | 4 | Yes | Yes |

Whilst useful, the pipelines mentioned above are all restricted to a small number of options at each stage, meaning that the user is unable to easily combine different sets of tools, which may be critical in certain settings. Here we present iRAP (an integrated RNA-seq Analysis Pipeline) that allows the user to choose the tool to be used on each stage of the analysis from amongst the many supported tools for mapping, quantification, differential expression analysis, and gene set enrichment analysis. iRAP performs the analysis from the raw reads up to the identification of differentially expressed genes or enriched set of genes with a single command. iRAP can also generate web reports to facilitate the inspection of the results produced at different stages of the analysis.

## 2 About iRAP

iRAP requires as input the FASTQ or BAM files containing the raw sequence reads, a reference genome (in FASTA format), gene model annotation (in GTF format) and a configuration file.

In the first step of the pipeline, the quality of the data is assessed and, optionally, the reads can be trimmed and those with “bad” quality filtered out.

The second step of the pipeline aligns the reads to the reference genome or transcriptome using one of the many available high-throughput sequencing mapping tools [4]. More than ten mappers, both splice aware and splice unaware are currently integrated in iRAP: Tophat1 [22], Tophat2 [9], Osa [7], Star [3], GSNAP [28], Bowtie1 [12], Bowtie2 [11], Smalt ^1^, BWA1 [13], BWA2 [14], GEM [17] and SOAPsplice [8]. The SAM/BAM files produced by the mappers are post-processed to ensure interoperability. This is necessary since the SAM/BAM files generated by some programs do not include all BAM fields required by downstream quantification tools (see appendix for more details).

The alignments generated are subsequently passed to one of four quantification tools to obtain a measure of expression over a biologically meaningful unit, such as exons, isoforms, or genes using Cufflinks [23, 24], FluxCapacitor [18], NURD [16], or HTSeq [2].

To identify differentially expressed genes, iRAP allows the user to select from one of the following tools: Cuffdiff1 [23], Cuffdiff2 [21], DESeq [1] or EdgeR [19]. Gene set enrichment (GSE) analysis can be performed using one of the many gene set analysis methods available in the Piano R package [26]. Finally, iRAP allows the user to quickly explore the results of the analysis through web browsable reports for each different stages of the analysis (QC, mapping, quantification, DE and GSE). Further details about the pipeline are provided in the Appendix.

## 3 Discussion

iRAP has a number of practical advantages. First it allows the practitioner to customise the analysis pipeline to the needs of a given project (e.g., question addressed, characteristics of the data, collaborators requests, etc) and change it easily as required. If there is uncertainty about the optimal pipeline setup, iRAP allows different options to be easily compared.

Additionally, iRAP is user friendly while maintaining a high level of flexibility. For instance, invoking iRAP as `irap conf=myexp.conf mapper=star quant_method=htseq de_method=deseq` would use Star as the mapper, HTSeq as the quantification tool and DESeq for differential expression. Information about the data is passed to iRAP in a configuration file (*myexp.conf* in the above example). Flexibility is provided by letting the user: i) select the tool to use on each stage of the pipeline; ii) stop and re-start the analysis pipeline at any stage; and iii) pass specific parameters to the tools used. Finally, iRAP can exploit multi-processor computers by defining the number of cores that can be used; the whole pipeline can also be executed in parallel on LINUX clusters that have a Load Sharing Facility (LSF). In the future we plan to integrate more methods in the pipeline to offer more alternatives and to address other biological questions (e.g., fusion detection, exon or isoform-level differential expression).

## Acknowledgments

We would like to thank iRAP’s users for their helpful and constructive comments.

## A iRAP details

iRAP requires as input the FASTQ files generated from an RNA-seq experiment, a reference genome (in FASTA format), gene model annotation (in GTF format) and an iRAP configuration file. The configuration file can contain options to run iRAP as well as information about the FASTQ files (read length, quality encoding, etc), experimental design (for differential expression analysis) and the location of the data (FASTQ files, genome and gene model).

The pipeline is controlled by a main GNU Make file, which supports a variety of programs for sequence alignment, quantification, and differential expression. The software also includes many utilities, custom scripts and programs that are used by the main GNU Make file to pre-process and post-process the many files generated when the pipeline is used.

iRAP avoids processing the same data set multiple times whenever possible. To do this, every time that iRAP is applied to the same input files, the software checks whether the files produced at each stage of the pipeline already exist and, if they do and are consistent, the existing files will be used instead of being recomputed. This is particularly useful since iRAP can be run up a particular stage (e.g., the end of the mapping) and stopped before re-starting the analysis.

### A.1 QC

Before aligning the reads, iRAP can optionally check the quality of the data by invoking FastQC^2^. If the user so wishes, reads can be trimmed and those with bad “quality” excluded from the analysis. The filtering is performed in four steps:

a) Base quality: the quality of the reads is assessed using the FASTX ^3^ toolkit. The reads are processed as follows: i) reads with an overall percentage (defined by the user) of the bases below some quality (default is 10) can be excluded; ii) bases at the ends of a read with a base quality below a user defined threshold can be trimmed; if the a read’s length decreases by more than 15%, it is discarded; iii) reads containing artifacts can be removed, where artifacts are defined using the `fastx artifacts filter`.
b) Contamination: reads that likely originate from organisms other than the one under study can be discarded. This is done by quickly aligning reads to the genomes of organisms that might be a source of contamination and those that map with a high degree of fidelity are discarded.
c) Uncalled bases: reads with uncalled bases (bases where the sequencer was unable to determine the base and assigned it an ‘N’) are excluded.
d) Unpaired reads: when reads are generated using a paired-end protocol, one of the pair can be lost in an earlier analysis step. These reads are placed into a singletons file, since most downstream analysis tools require paired end-reads to be separated into two FASTQ files, where the pairs are listed in the same order. Unpaired reads are assigned to a third file.

Lastly, the filtered reads are processed using FastQC to generate an HTML report of the quality of the data.

### A.2 Mapping

Aligning the (quality filtered) reads to the reference genome is a key step in the analysis of RNA-seq data. iRAP allows the user to select from one of the following mappers: Tophat 1 [22], Tophat 2 [9], Osa [7], Star [3], GSNAP [28], Bowtie2 [11], Bowtie1 [12], Smalt ^4^, BWA1 [13], BWA2 [14], GEM [17] and SOAPsplice [8]. Other mappers can also be included if they output aligned reads in the SAM or BAM format. Most HTS mappers require the reference genome to be indexed [4]. This step is automatically performed by iRAP for the selected mapper. Moreover, some HTS mappers also require a file with splice junctions to be inputted. Again, iRAP automatically generates this file using the reference and annotation files provided as input.

The SAM/BAM files produced by the mappers are post-processed in two steps. First, to ensure interoperability, the SAM/BAM files are processed to include, if necessary, fields that the mappers failed to generate, but that may be required by downstream quantification tools (see Table 2 and Table 3).

**Table 2:** SAM/BAM fields used by different quantification methods. The NH field contains information about the number of alignments associated with a read; the XS:A field contains the splice strand orientation; and Splicing indicates whether splicing is reported in the cigar string.

| Method | NH | XS:A | Splicing (Cigar) |
| --- | --- | --- | --- |
| HTSeq | Yes | No | No |
| Cufflinks 1 and 2 | Yes | Yes | Yes |
| Flux-Capacitor | Yes | Yes | Yes |
| NURD | No | No | No |

**Table 3:** SAM/BAM fields generated by the different mappers (Yes - generated, No - not generated).

| Mappers | NH | XS:A | Splicing (cigar) |
| --- | --- | --- | --- |
| Tophat 1 | Yes | Yes | Yes |
| Tophat 2 | Yes | Yes | Yes |
| Bowtie 1 | No | No | No |
| Bowtie 2 | No | No | No |
| Smalt | No | No | No |
| Gsnap | Yes | Yes | Yes |
| Soapsplice | No | No | Yes |
| BWA 1 | No | No | No |
| BWA 2 | No | No | No |
| Gem | Yes | No | No |
| OSA | Yes | Yes | Yes |
| Star | Yes | Yes | Yes |

For each input library, the output of the mapping step is three files: a) a BAM file with the alignments sorted by chromosomal coordinates; b) a BAM file with the alignments sorted by read name; c) a BAM index for the file sorted by chromosomal coordinates. These files are generated using SamTools [15]. Obviously, generating two BAM files for the same data set is wasteful but this is necessary since some downstream tools require alignments to be sorted in different ways.

### A.3 Quantification

After mapping, the next task is to summarise and aggregate reads over a biologically meaningful unit, such as exons, isoforms, or genes. iRAP supports several quantification tools: HTSeq ^5^, Cufflinks 1 [23], Cufflinks 2 [24], NURD [16], and Flux-capacitor [18]. Subsequently, iRAP processes the output files to generate a matrix with the number of reads per biological unit and per BAM files. Depending upon the quantification tool selected, iRAP produces a TSV file with the read count matrix per gene, exon or isoform.

### A.4 Differential Expression

iRAP allows the user to identify differentially expressed genes using one of the following tools: Cuffdiff 1 [23], Cuffdiff 2 [21], DESeq [1] or EdgeR [19]. Current strategies for integrated DE analysis are limited to simple experiment designs, such as pairwise comparisons. For more complex designs, such as paired samples or time course experiments, the user needs to get the data and apply an appropriate alternative tool.

### A.5 Gene set enrichment analysis

Gene set enrichment (GSE) analysis can be performed using the (adjusted) p-values and fold-change computed for each gene during the differential expression analysis. iRAP performs GSE analysis using the Piano R package [26] which provides support for several statistical methods (e.g, fisher, stouffer,…). The output of a GSE analysis will be a TSV file with gene set statistics and p-values.

### A.6 Web reports

iRAP allows the user to quickly explore the results of the analysis through web browsable reports for each different stages of the analysis (QC, mapping, quantification, DE and GSE). For each stage of the analysis one or several web pages are generated with tables and plots.

## B Installation and usage

iRAP integrates many third party tools, all of which need to be installed. To facilitate this task, iRAP includes an install script to compile and install all third-party methods and tools in a user defined folder. A Virtual Machine is also provided with iRAP and all dependencies pre-installed. Although the pipeline should be portable to any POSIX-compliant operating system, several third party methods only run on Linux. As a result, iRAP can only be installed on a Linux operating system with development libraries installed — these will be needed to compile and install Perl, R, Ruby and other utilities in a folder defined by the user. No administration access to the operating system is required.

As mentioned above, iRAP requires as input FASTQ files containing the reads, a reference genome (in FASTA format), gene model annotation (in GTF format) and a configuration file. Then, assuming that the configuration file is called `myexp.conf`, the pipeline can be started by invoking iRAP: `irap conf=myexp.conf`. The specific combination of tools required can be easily specified from the command line. For instance, the Tuxedo pipeline can be defined as `irap conf=myexp.conf mapper=tophat1 quant_method=cufflinks1 de_method=cuffdiff1`. Alternatively, the combination of Star, HTSeq and DESeq can be specified as `irap conf=myexp.conf mapper=star quant_method=htseq de_method=deseq`.

It is possible to run iRAP only up to some stage of the pipeline. For instance, `irap conf=myexp.conf mapping` will run the pipeline up to the mapping stage.

Processing RNA-sequencing data is often computationally demanding both in terms of memory requirements and in CPU time. To reduce the execution time iRAP can exploit the multiple processors and cores available in a modern computer as well as utilising a Linux cluster. Furthermore, iRAP exploits the Platform LSF job scheduler by splitting the analysis into multiple jobs with the aim of reducing the time to produce the results. From the user perspective, this can be achieved without changing the parameters, input files or configuration file.

Finally, it is important to keep track of the software used and the respective versions. iRAP provides this information alongside a bibliographic citation of the tools used (if available) via a simple command (e.g.,`irap conf=myexp.conf show_citations`). Furthermore, iRAP provides all commands executed along with the input and output files providing the user with a log of all commands executed.

## Footnotes

1 Smalt: http://www.sanger.ac.uk/resources/software/smalt

2 FastQC: http://www.bioinformatics.babraham.ac.uk/projects/fastqc

3 FASTX: http://hannonlab.cshl.edu/fastx_toolkit/

4 Smalt: http://www.sanger.ac.uk/resources/software/smalt/

5 HTSeq: http://www-huber.embl.de/users/anders/HTSeq/

## References

1. S. Anders and W. Huber. Differential expression analysis for sequence count data. Genome Biol, 11(10):R106, 2010.

2. Simon Anders, Paul Theodor Pyl, and Wolfgang Huber. HTSeq–A Python framework to work with high-throughput sequencing data. bioRxiv, 2014.

3. Alexander Dobin, Carrie A Davis, Felix Schlesinger, Jorg Drenkow, Chris Zaleski, Sonali Jha, Philippe Batut, Mark Chaisson, and Thomas R Gingeras. STAR: ultrafast universal RNA-seq aligner. Bioinformatics, 29(1):15–21, 2013.

4. Nuno A. Fonseca, Johan Rung, Alvis Brazma, and John Marioni. Tools for mapping high-throughput sequencing data. Bioinformatics, 28(24):3169– 3177, December 2012.

5. Jeremy Goecks, Anton Nekrutenko, James Taylor, T Galaxy Team, et al. Galaxy: a comprehensive approach for supporting accessible, reproducible, and transparent computational research in the life sciences. Genome Biol, 11(8):R86, 2010.

6. Ângela Gonçalves, Andrew Tikhonov, Alvis Brazma, and Misha Kapushesky. A pipeline for rna-seq data processing and quality assessment. Bioinformatics, 27(6):867–869, 2011.

7. J. Hu, H. Ge, M. Newman, and K. Liu. OSA: a fast and accurate alignment tool for RNA-Seq. Bioinformatics, 28(14):1933–1934, 2012.

8. Songbo Huang, Jinbo Zhang, Ruiqiang Li, Wenqian Zhang, Zengquan He, Tak-Wah Lam, Zhiyu Peng, and Siu-Ming Yiu. SOAPsplice: genome-wide ab initio detection of splice junctions from RNA-Seq data. Frontiers in Genetics, 2(0), 2011.

9. Daehwan Kim, Geo Pertea, Cole Trapnell, Harold Pimentel, Ryan Kelley, and Steven L Salzberg. Tophat2: accurate alignment of transcriptomes in the presence of insertions, deletions and gene fusions. Genome biology, 14(4):R36, 2013.

10. David G Knowles, Maik Röder, Angelika Merkel, and Roderic Guigó. Grape RNA-Seq analysis pipeline environment. Bioinformatics, 29(5):614–621, 2013.

11. B. Langmead and S.L. Salzberg. Fast gapped-read alignment with Bowtie 2. Nature methods, 9(4):357–359, 2012.

12. B. Langmead, C. Trapnell, M. Pop, S.L. Salzberg, et al. Ultrafast and memory-efficient alignment of short DNA sequences to the human genome. Genome Biol, 10(3):R25, 2009.

13. H. Li and R. Durbin. Fast and accurate short read alignment with burrows– wheeler transform. Bioinformatics, 25(14):1754–1760, 2009.

14. H. Li and R. Durbin. Fast and accurate long-read alignment with burrows–wheeler transform. Bioinformatics, 26(5):589–595, 2010.

15. H. Li, B. Handsaker, A. Wysoker, T. Fennell, J. Ruan, N. Homer, G. Marth, G. Abecasis, R. Durbin, et al. The sequence alignment/map format and SAMtools. Bioinformatics, 25(16):2078, 2009.

16. Xinyun Ma and Xuegong Zhang. NURD: an implementation of a new method to estimate isoform expression from non-uniform RNA-seq data. BMC Bioinformatics, 14(1):220, 2013.

17. S. Marco-Sola, M. Sammeth, R. Guigó, and P. Ribeca. The GEM mapper: fast, accurate and versatile alignment by filtration. Nature Methods, 9(12):1185–1188, 2012.

18. Stephen B Montgomery, Micha Sammeth, Maria Gutierrez-Arcelus, Radoslaw P Lach, Catherine Ingle, James Nisbett, Roderic Guigo, and Emmanouil T Dermitzakis. Transcriptome genetics using second generation sequencing in a caucasian population. Nature, 464(7289):773–777, 2010.

19. MD Robinson, DJ McCarthy, and GK Smyth. edgeR: a Bioconductor package for differential expression analysis of digital gene expression data. Bioinformatics, 26:139–140, 2010.

20. Wandaliz Torres-García, Siyuan Zheng, Andrey Sivachenko, Rahulsimham Vegesna, Qianghu Wang, Rong Yao, Michael F Berger, John N Weinstein, Gad Getz, and Roel GW Verhaak. Prada: Pipeline for rna sequencing data analysis. Bioinformatics, page btu169, 2014.

21. C. Trapnell, D.G. Hendrickson, M. Sauvageau, L. Goff, J.L. Rinn, and L. Pachter. Differential analysis of gene regulation at transcript resolution with rna-seq. Nature Biotechnology, 2012.

22. C. Trapnell, L. Pachter, and S.L. Salzberg. TopHat: discovering splice junctions with RNA-Seq. Bioinformatics, 25(9):1105–1111, 2009.

23. C. Trapnell, B.A. Williams, G. Pertea, A. Mortazavi, G. Kwan, M.J. Van Baren, S.L. Salzberg, B.J. Wold, and L. Pachter. Transcript assembly and quantification by RNA-Seq reveals unannotated transcripts and iso-form switching during cell differentiation. Nature biotechnology, 28(5):511– 515, 2010.

24. Cole Trapnell, Adam Roberts, Loyal Goff, Geo Pertea, Daehwan Kim, David R Kelley, Harold Pimentel, Steven L Salzberg, John L Rinn, and Lior Pachter. Differential gene and transcript expression analysis of RNA-seq experiments with TopHat and Cufflinks. nature protocols, 7(3):562–578, 2012.

25. Ernest Turro, Shu-Yi Su, Ângela Gonçalves, LJ Coin, Sylvia Richardson, and Alex Lewin. Haplotype and isoform specific expression estimation using multi-mapping RNA-seq reads. Genome Biol, 12(2):R13, 2011.

26. Leif Väremo, Jens Nielsen, and Intawat Nookaew. Enriching the gene set analysis of genome-wide data by incorporating directionality of gene expression and combining statistical hypotheses and methods. Nucleic acids research, 41(8):4378–4391, 2013.

27. Ying Wang, Gaurang Mehta, Rajiv Mayani, Jingxi Lu, Tade Souaiaia, Yangho Chen, Andrew Clark, Hee Jae Yoon, Lin Wan, Oleg V Evgrafov, et al. Rseqflow: workflows for rna-seq data analysis. Bioinformatics, 27(18):2598–2600, 2011.

28. TD Wu and S Nacu. Fast and SNP-tolerant detection of complex variants and splicing in short reads. Bioinformatics, 26:873–881, 2010.

